# Cell type-specific complement expression from healthy and diseased retinae

**DOI:** 10.1101/413088

**Authors:** Diana Pauly, Nicole Schäfer, Felix Grassmann, Anna M. Pfaller, Tobias Straub, Bernhard H. F. Weber, Stefanie M. Hauck, Antje Grosche

## Abstract

Retinal degeneration is associated with complement system activation, but retinal sources of complement are unknown. Here, we describe the human and murine complement transcriptomes of Müller cells, microglia/macrophages, vascular cells, neurons and retinal pigment epithelium (RPE) in health and disease. All cell populations expressed *c1s, c3, cfb, cfp, cfh* and *cfi*. Murine Müller cells contributed the highest amount of complement activators (*c1s, c3, cfb*). RPE mainly expressed *cfh,* while *cfi* and *cfp* transcripts were most abundant in neurons. The main complement negative regulator in the human retina was *cfi*, while *cfh* dominated in the murine retina. Importantly, the expression of *c1s, cfb, cfp, cfi* increased and that of *cfh* decreased with aging. Impaired photoreceptor recycling led to an enhanced *c3* expression in RPE and to a reduced *cfi* expression in microglia/macrophages. Expression of complement components was massively upregulated after transient retinal ischemia in murine microglia, Müller cells and RPE. The individual signature of complement expression in distinct murine and human retinal cell types indicates a local, well-orchestrated regulation of the complement system in both species.

## Introduction

The retina, as the light sensitive part of the eye, consists of more than 40 different cell types, which cooperate to capture, process, and transmit visual signals ^1–4^. Over the past, our understanding of the functional orchestration of these processes in healthy and diseased retinae and its supporting tissues like the retinal pigment epithelium (RPE), retinal and choriocapillary vessels has grown considerably, including the characterization of the transcriptomic landscape of the retina ^2,5^. However, most of these studies were performed based on RNA from whole retinae losing the information from which retinal cell type the detected RNA originated ^2,5,6^. Droplet-based single-cell RNA sequencing and microarray analysis allowed studying the molecular differences between different subsets of retinal ganglion cells ^3^, bipolar cells ^4^ and a description of the Müller cell transcriptome ^7^. However, these studies have not been able to determine how major retinal cell types (Müller cells, vascular cells, microglia/macrophages, retinal neurons and RPE cells) shape the overall retinal transcriptome. An unresolved obstacle of single-cell RNAseq is that these data can be intrinsically noisy and result in low detection efficiencies for certain transcripts due to unevenness of sequencing depth across the global transcriptome ^8^.

Mutations in over 300 genes (https://sph.uth.edu/retnet/) are known to cause visual impairment because of retinal dysfunction. To study the mechanisms of retinal degeneration in depth, the role of retinal cell populations expressing pathology-associated genes and their contribution to retinal homeostasis is required. Polymorphisms in genes of the complement system, a pathway of the innate immune system, are associated with a number of retinal diseases. In glaucoma, mainly complement components of the classical ^9^ and terminal ^10^ pathway are affected, while age-related macular degeneration (AMD) ^11^ and diabetic rethinopathy ^12,13^ have essentially been correlated to polymorphisms of the alternative pathway. The complement pathway consists of over 40 proteins interacting in enzyme cascades that result in protein cleavage products. These fragments act as anaphylatoxins (e.g. C3a, C5a) which modulate cell functions via specific receptors. Moreover, opsonins are formed (e.g. C3b) that bind to cell surfaces and mark them for phagocytosis or that become a part of the membrane attack complex leading to cell activation or lysis ^14^. The classical pathway is activated by binding of the C1-complex to antibodies. Subsequently, the activating protease C1s within this complex initiates cleavage of downstream complement components forming the classical C3-convertase. This enzyme complex cleaves C3 into C3a and C3b – the latter being the common central step of all complement pathways. However, the C3-convertase of the alternative pathway consists of cleavage products of complement components C3 and complement factor B (CFB). This enzyme complex is strictly regulated by the interacting inhibitory complement factors H (CFH) and I (CFI). In contrast, the antagonistic complement factor P (CFP) stabilizes the alternative C3-convertase. The terminal complement pathway is characterized by formation of the C5-convertase and the membrane attack complex. A growing body of evidence suggests that complement components are locally expressed in the retina and RPE ^2,15–23^ and that activation is independent of the systemic complement cascades. The latter are produced in liver hepatocytes and distributed via the bloodstream. The extrahepatic, retinal complement system may facilitate a rapid response to microbial invasion and disposal of altered cells despite an intact blood-retina barrier. To date, only microglia ^23,24^ and RPE ^15,16,25^ cells have been identified as source of complement components in the retina. The remaining cell types have not been characterized so far.

Complement expression and deposition change in the aging ^26–28^ and diseased retina ^29–34^. Interestingly, formation of the membrane attack complex is enhanced with advancing age as demonstrated in human postmortem eyes ^35^. Moreover, complement components are present in extracellular deposits (termed “drusen”) which are the hallmark of AMD ^36,37^. Consequently, it is tempting to speculate that a source for retinal complement activation during aging could originate from the retina itself as animal studies showed an increased expression of *c1q, c3, c4* and *cfb* in retinae of older mice/rats ^26,27,38^. Apart from aging, complement upregulation was also shown for hereditary diseases like retinitis pigmentosa in dogs ^39^ as well as in a mouse model for Stargardt disease ^29,31^. Enhanced expression of the central complement component C3 indicated changes in the complement homeostasis towards higher complement activity in all of these hereditary diseases ^29,31,39^. An early involvement of the complement pathways and changes in complement expression were also suggested in non-congenital retinal diseases such as diabetic retinopathy ^40^ and glaucoma ^41^. Both conditions are associated with transient ischemic tissue damage and a pronounced concomitant expression of complement components has been demonstrated in several animal studies ^32–34^. In spite of a clear indication of a fundamental role of the complement system in retinal physiology and pathophysiology, there is a lack of knowledge about which retinal cell population is involved in shaping complement homeostasis in the healthy, aging or diseased retina.

Here, we profile the complement expression in five cellular fractions, each highly enriched for either Müller cells, vascular cells, microglia/macrophages, neurons or RPE cells of murine and human retinae, respectively. We observed a characteristic contribution of complement transcripts from each distinct retinal cell population to the retinal complement homeostasis. There were remarkable differences in complement transcription patterns between human and mouse retinal cell types, between pigmented and albino mice, in aging mice as well as in mouse models of acute or chronic retinal degeneration. Based on our data, we suggest *cfi* as the major complement inhibitor in the human and *cfh* in the mouse retina. A common mechanism during retinal stress appears to be the downregulation of *cfh* expression in mouse models of retinal degeneration. Apart from this, the cell type-specific changes in complement expression differed in aging and in ABCA4^-/-^ mice (model for Stargardt disease) as well as in acute retinal degeneration induced by transient ischemia, which indicated a stress-dependent cell type-specific modulation of the complement homeostasis in the retina.

## Results

### Major contribution of neurons, Müller cells and RPE to the retinal transcriptome

Investigating the role of five different retinal cell populations required their enrichment from the murine and human retina **(Fig 1A)**. We purified Müller cells, microglia/macrophages (in the following termed microglia, although contaminations with macrophages cannot be excluded), vascular cells and retinal neurons by immunomagnetic cell separation from albino BALB/c, pigmented C57BL/6 mice and human retinae ^42^. RPE was scratched from eyecups after the retina had been removed. Accordingly, RPE samples always also contained cells from the choroid. All five cell populations were characterized by testing the expression of specific marker genes **(Fig 1B)**. Gene expression of the glial marker glutamine synthetase (*glul*) was mainly enriched in the Müller cell fraction. The microglia marker gene integrin alpha M (*cd11b*) was preferentially expressed in the microglia population, the platelet endothelial cell adhesion molecule (*pecam*) mRNA characterized the purified vascular cells and the mRNA of the neural leucine zipper (*nrl*) was a marker gene for photoreceptors. The retinoid isomerohydrolase (*rpe65*) was specifically detected in the RPE **(Fig 1B)**. The efficiency of enrichment of Müller cells, vascular cells and photoreceptors was similar in 8, 16 and 24 week old albino mice. Of note, we found a rise of *cd11b*-expression in microglia and reduction of *rpe65* transcript levels in RPE with increasing age **(Fig 1B)**.

**Figure 1.**
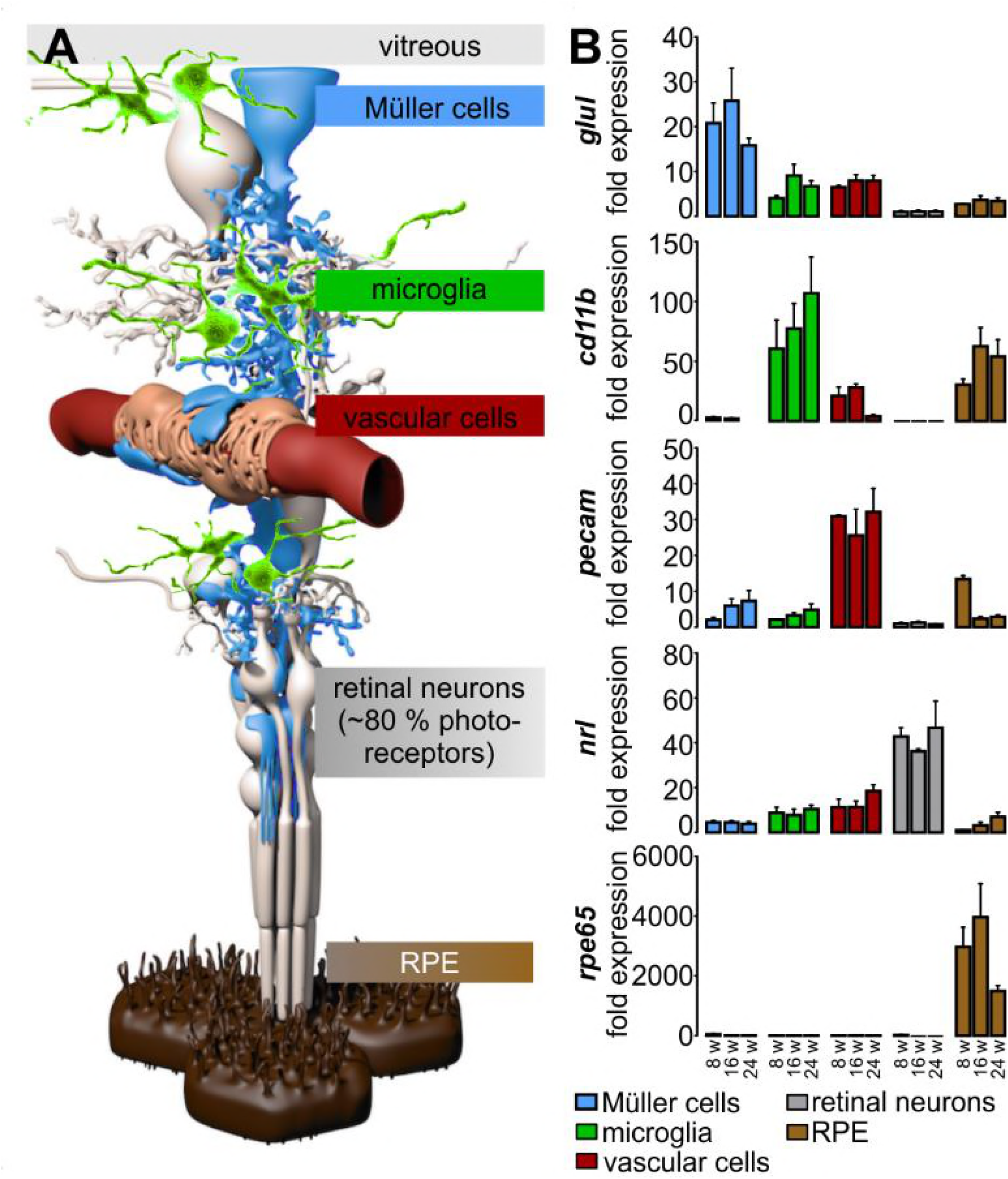
Validation of enrichment of different retinal cell types. A Schematic view of distinct retinal cell types. Müller cells (blue), the central glia cells of the retina, are in direct contact with the vitreous and various retinal cell types: microglia (green), vascular cells (red) and neurons (grey). 80% of the retinal neurons are light-responsive photoreceptors that are supported by retinal pigment epithelial cells (RPE, brown). B Murine retinal cell populations were enriched by immunomagnetic cell separation and characterized by qRT-PCR using specific markers: *Glul* is a marker for the Müller cell fraction. Microglia (and putatively co-enriched macrophages) specifically express *cd11b.* Vascular cell enrichment was proven by strong expression of *pecam*. Retinal neurons were characterized by an enhanced detection of the photoreceptor-specific *nrl* mRNA compared to the other cell populations. *Rpe65* was exclusively expressed in RPE/choroid. Exemplarily shown mean values ±EM for cell preparations from BALB/c mice 8, 16 and 24 weeks of age (n=4 – 6 for each age).

Expectedly, transcripts for ionized calcium binding adaptor molecule 1 (*iba1*) and intercellular adhesion molecule 1 (*icam-1*) were detected in RPE samples from pigmented C57BL/6 and albino mice indicative of a contamination with macrophages residing in choroidal vessels and vascular cells **(Fig EV1)**, respectively. While tests to deplete RPE of pigmented C57BL/6 mice from contaminating cells were successful **(Fig EV1)**, the protocol was not feasible for RPE from albino BALB/c mice given the considerably lower yields in cell numbers compared to that from eyecups of pigmented mice. Consequently, for interpreting the complement component expression data we had to consider minimal macrophage and vascular cell contaminations in the mixed RPE/choroid samples **(Fig 2, 3, EV4, 4)**.

**Figure 2.**
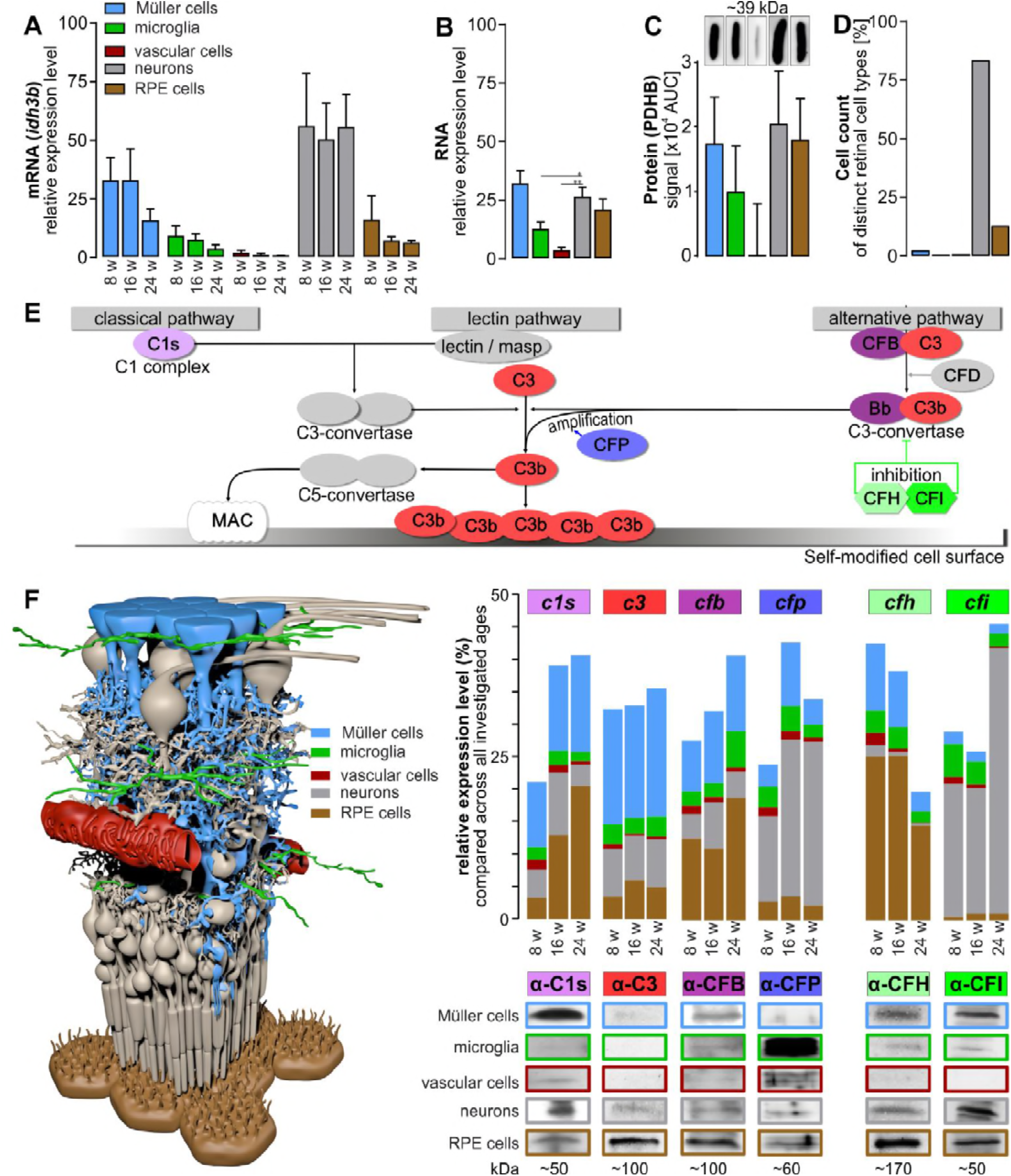
Contribution of different retinal cell types to the retinal architecture, expressome and complement homeostasis in aging albino wild type mice. A The mRNA expression levels of the newly identified and validated housekeeping gene *idh3b* was determined in samples of the five different retinal cell populations from retinae of 8, 16 and 24 week old albino mice without adjusting the cell type-specific amount of RNA input. This allows a rough estimate of the overall relative contribution to of each cell type to the retinal transcriptome. Bars represent mean values ± SEM (n=4 – 6). B Similar to (A), the total RNA amount of the five different retinal cell types enriched from 8 to 16 weeks old albino mice was investigated using Agilent RNA 6000 picochip analysis. Note that Müller cells, neurons and RPE/choroid cells yielded comparable amounts of mRNA. Bars represent mean values ± SEM (n=5 – 8). *P<0.05, **P<0.01, Mann-Whitney U-test. C Quantification of PDHB protein expression in Western Blots suggested comparable PDHB protein levels in the Müller cell, retinal neuron and RPE/choroidal cell population in the murine retina. Western Blots were performed from five different cell preparations purified from 4 – 6 albino mice. D Previously published ^43^ and own counted relative retinal cell numbers in a mouse retina. E Schematic overview of the complement system. The pathway can be activated via three different mechanism and is amplified by an amplification loop. Note that the present study focussed on the characterization of expression profiles of the coloured complement components C1s, C3, CFB, CFP, CFH and CFI. F Expression levels of complement components C1s, C3, CFB, CFP, CFH and CFI were determined in 8, 16 and 24 week old albino mice at mRNA level (bar graphs) and in 24 week old mice at protein level (Western blot data). Determination of the overall proportional contribution of each cell population to the local complement homeostasis was performed by analysing the total yield of mRNA or protein derived from the respective cell population from two mouse retinae. Intraretinal complement expression is dominated by Müller cells and neurons, while expression changes in vascular cells should have minor effects.

**Figure 3.**
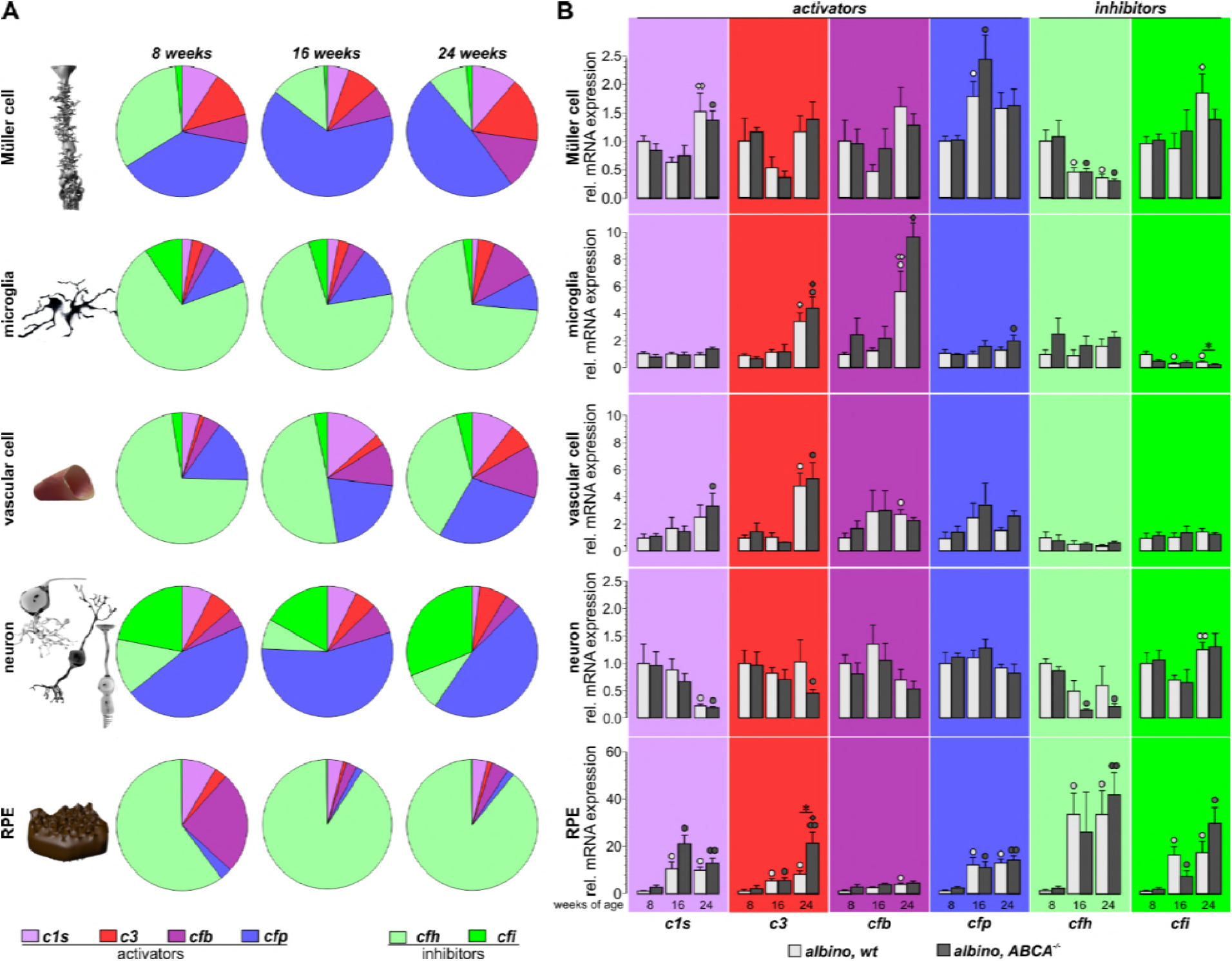
Comparison of complement component expression between retinal cell types of aging albino wild type and ABCA4-/- mice. A Expression of complement components was determined by qRT-PCR. Diagrams represent the relative amount of transcripts per cell (normalized to the *idh3b* housekeeper expression) of the different complement components in the respective cell type enriched from mice at the indicated age. Note the high expression level of inhibitory complement factors in RPE/choroid samples as well as in microglial and vascular cells, while complement activating genes appear to dominate in Müller cells and neurons. Data were collected from 4 – 6 wild type albino mice (numbers are given in **Table S2**). B Complement expression analysis by qRT-PCR was performed on enriched retinal cell types from 8, 16 and 24 week old wild type and ABCA4^-/-^ mice. ABCA4^-/-^ mice presented with a significant enhanced expression of *c3* in RPE cells and a decreased expression of *cfi* in microglia cells compared to wild type controls. Age-dependent changes in complement expression were similar in both mouse strains to the greatest extent. Bars represent mean values ± SEM of cells purified from 4 – 6 animals. Mann-Whitney U-testing was performed on all data. *P<0.05. Circle, significant difference compared to the expression level at 8 weeks of age; diamond, significant difference compared to the expression level at 16 weeks of age. °/◊P<0.05; ° °/◊ ◊P<0.01

**Figure 4.**
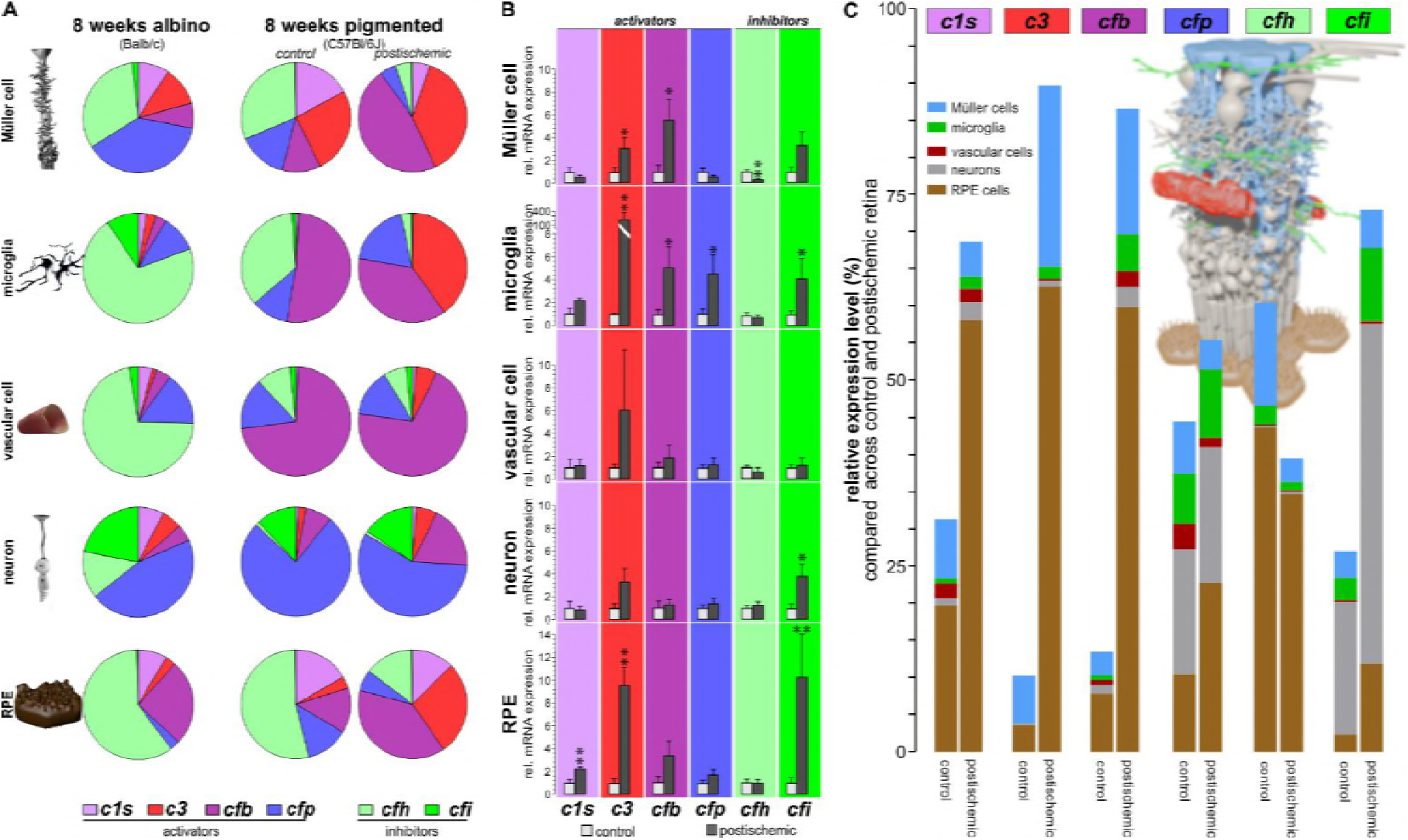
Transient ischemic stress results in cell type-specific upregulation of transcripts from activating complement components and downregulation of cfh. A The relative amount of complement transcripts per retinal cell type indicated that complement activating transcripts are more abundant in pigmented mice than in albino mice in which transcripts of complement inhibitors dominate at the same age. Note the strong relative upregulation and the resulting shift towards transcripts from complement activators 24 hours after transient ischemic retinal stress in all retinal cell types of C57BL/6 mice (numbers are given in **Table S3**). B Major changes of local complement expression (normalized to the house keeper) were detected by qRT-PCR 24 hours after transient ischemia. The most pronounced upregulation of complement activators was found in Müller cells (*c3, cfb*), microglia (*c3, cfb, cfp*) and RPE (*c1s, c3*). *Cfh* was down-regulated in Müller cells, while *cfi* was up-regulated in all investigated cell types. Significantly different expression as compared to that cells from healthy control eyes: *P<0.05, **P<0.01, Mann-Whitney U-test. C Complement transcript contribution of the different retinal cell populations (no normalization to the housekeeper and no adjustment of RNA input) indicated a pro-inflammatory milieu in the postischemic retina. Mainly Müller cells, microglia and RPE cells contributed to the changed complement homeostasis in postischemic retinae. A – C Data were collected from 3 – 5 animals.

To normalize expression, we needed to identify novel housekeeping genes with suitable expression across all retinal cell populations. Commonly used housekeeping genes such as β-actin (*actb*) or glycerinaldehyd-3-phosphat-dehydrogenase (*gapdh*) showed surprisingly high transcriptional and even more pronouncedly translational variability when comparing the different cell types (**Fig EV2A-D**). Using data from an RNA sequencing approach **(Fig EV2E, H)**, validation by quantitative RT-PCR **(Fig EV2F, I)** and finally also at protein level applying quantitative mass-spectrometry **(Fig EV2G, J)**, we found pyruvate dehydrogenase E1 component subunit beta (*pdhb*) **(Fig EV2E-G)** and isocitrate dehydrogenase 3 (NAD+) beta (*idh3b*) **(Fig EV2H-J)** as being expressed at similar levels in all five cell populations we investigated at mRNA and protein level if related to whole RNA or protein input amounts, respectively. Accordingly, we considered these genes more suitable as reference housekeeping genes **(Fig 2, 3, EV4, 4)**.

Using *idh3b* to determine the proportional contribution of a distinct cell population to the total retinal transcriptome in albino mice, we identified the neuronal fraction (60%) as the cell population expressing highest levels of transcripts **(Fig 2A).** Müller cells contributed 25% of the retinal *idh3b* transcript. RPE (9%), vascular cells (1%) and microglia (5%) together expressed less *idh3b* than Müller cells or neurons alone, indicating low cell numbers and/or low transcription activity of the former cell types in the mouse retina. Quantifying whole RNA by RNA picochip analysis, we found similar RNA amounts in Müller cells (33%), neurons (26%) and the RPE/choroid fraction (22%) per mouse **(Fig 2B)**. In line with this, comparing the protein levels of PDHB as a second indicator for cell contributions in the mouse eye, we found comparable expression signals in Müller cells, neurons and RPE cells **(Fig 2C)**, while microglia and vascular cells showed lower PDHB signals. According to previously published and own retinal cell counts, neurons contribute up to 85% of the total retinal and associated RPE cell population, RPE cells 13%, Müller cells 2% and microglia as well as vascular cells below 1% in the mouse **(Fig 2D)** ^43^. Given our results regarding mRNA **(Fig 2A)**, RNA **(Fig 2B)** and protein **(Fig 2C)** expression levels, these numbers underestimated the role of Müller cells and microglia cells regarding their role in shaping the retinal microenvironment by gene expression. Müller cell numbers only contribute 2% of total retinal cell numbers, but they contribute up to 25% of the total mRNA content in the retina based on housekeeping expression levels. To our knowledge, this is the first study providing insight into the proportional transcriptional activities of the five major cell populations in the retina.

### Retinal cell populations contribute differently to the intraretinal complement homeostasis

To address our working hypothesis, which assumes that different retinal cell types shape the intraretinal complement homeostasis by expressing specific sets of components from the complement pathway, we analysed the expression patterns of six complement genes across distinct cell types **(Fig 2E, F)**. We chose the complement activating genes of the classical (*c1s*) and of the alternative pathway (*cfb*), which were of interest as they have been associated with retinal degeneration ^11,15,44^. C3 is the central complement component. Its cleavage mediates the main complement functions, rendering it an ideal candidate for the study. We further analysed complement regulators of the amplification loop that contribute to all three activation pathways. The amplification loop is stabilised by CFP and inhibited by CFH and CFI **(Fig 2E)**. Implementing our novel cell type-specific expression approach, we detected transcripts of all investigated complement activators and regulators in every retinal cell type tested **(Fig 2F)**. Importantly, in order to characterize the contribution of whole cell populations to the overall retinal expression level of a respective complement component, no normalization to the housekeeping expression was performed. The Müller cell population contributed the highest amount of transcripts of complement activators, expressing 34 – 47% of *c1s*, 53 – 56% of *c3* and 28 – 34% of *cfb* retinal transcripts, depending on the mouse age of 8 to 24 weeks **(Fig 2F)**. Retinal neurons dominated the expression of the complement regulators *cfi* and *cfp*, while 59 – 74% of the *cfh* transcripts were detected in the RPE. Of note, despite the relatively low number and transcript amount of microglia **(Fig 2A – D)**, the resident immune cells of the retina contributed proportionally more *cfh* mRNA and a similar amount of *cfb* transcripts to the retinal complement homeostasis compared to, e.g., retinal neurons **(Fig 2F)**.

Typically, complement components are secreted, circulate in the blood stream and act on modified surfaces. In Western blot experiments, we found that the complement activators C1s, CFB and CFP were detectable in all enriched and washed murine cell populations, albeit at different levels **(Fig 2F)**. The main complement component C3 could be detected in RPE, Müller cells and neurons, while the complement inhibitors CFH and CFI were present in all cell types except for vascular cells. The complement component transcript level as determined by qRT-PCR **(Fig 2F)** did largely, but not completely match the pattern of complement proteins detected in the different, washed retinal cell populations **(Fig 2F)**. High transcript level of *c1s* were found in Müller cells **(Fig 2F)** matching the prominent C1s protein detection **(Fig 2F)**. Depending on the age, neurons expressed 39 – 40 % of retinal *cfp* mRNA **(Fig 2F)**, but the highest concentration of CFP protein was associated with microglia cells and not with neurons in the albino retinae **(Fig 2F)**. This may imply a spatial separation of complement component transcription and complement component accumulation at the protein level within the retina.

Collectively, these experiments showed: (i) complement components are locally expressed by the five different retinal cell populations investigated. (ii) Given the cell-specific ratios of activating and inhibiting complement component expression levels, each cell entity may have its specific role maintaining the retinal complement homeostasis. (iii) Cell populations present in the retina at low numbers such as microglia and Müller cells can have a major impact on retinal complement activity.

### Age-dependent changes in the complement expression of different retinal cell populations from albino mice

Albino BALB/c mice showed age-dependent complement expression patterns in the different retinal cell populations. The sum over all cell populations *c1s, cfb, cfp* and *cfi* transcripts increased over time, while transcript expression of *c3* was stable in 8 to 24 week old mice and that of *cfh* decreased with aging **(Fig 2F)**.

RPE cells of 16 to 24 week old BALB/c mice expressed 4 – 6 times more *c1s* than those of 8 week old mice. Given that besides Müller cells and neurons, RPE cells were the main *c1s* producers in the retina, this age-dependent rise in *c1s* expression in the RPE resulted in a doubling of overall retinal *c1s* mRNA in 24 week old compared to 8 week young mice **(Fig 2F)**.

The central complement component *c3* was mainly transcribed in Müller cells and the ratio of the producing cell entities stayed constant in aging mice. Only in vascular cells the *c3* mRNA was reduced by 50% in 24 week old animals **(Fig 2F)**. However, this may barely affect the retinal complement homeostasis, since the vascular cell population is small and thus contributes only little to the overall retinal *c3* expression **(Fig 2A – D)**.

Transcripts of the alternative pathway activator *cfb* were primarily produced by Müller cells and the RPE **(Fig 2F)**. *Cfb* expression increased in these cell populations in 24 week old mice compared to 8 week controls, which resulted in 1.5-fold higher overall retinal *cfb* level. An interesting change was observed when comparing *cfb* transcription in neurons and microglia. The neuronal population expressed a constant amount of *cfb* in 8 and 24 week old albino mice, but in the microglial population, the *cfb* transcript levels increased 2.6-fold in 24 week compared to 8 week old animals. This resulted in a higher *cfb* mRNA contribution of the small microglial population in older animals (14%) compared to 10% *cfb* transcript coming from the largest retinal cell population, the neurons, in aged mice **(Fig 2)**.

The transcript of the only known positive regulator, *cfp*, was mainly synthetized by neurons (55%) in retinae of juvenile mice. The expression ratio was further shifted towards the neuronal population (73%) in 24 week old animals, as *cfp* mRNA expression decreased in Müller and RPE cell populations with aging and rose in retinal neurons. Interestingly, *cfp* was the only transcript, which showed the highest transcript amount at 16 weeks of age compared to 8 and 24 weeks of age, which was caused by a temporarily higher *cfp* expression by Müller cells and a moderately enhanced expression in RPE cells at that age **(Fig 2F).**

CFH is a CFP antagonist and the main negative regulator of the complement system. The overall retinal *cfh* transcript level was reduced by 50% in retinae of 24 week old as compared to 8 week old mice. Consistently, all investigated cell populations decreased *cfh* transcript expression during aging with more pronounced effects found in microglial and RPE cell populations. Of note, 73% of the retinal *cfh* transcript is produced by RPE cells in the 24 week old retinal microenviroment of albino mice **(Fig 2F)**.

CFH is an important cofactor in complement inhibition for CFI. Interestingly, up to 90% of the *cfi* transcript was transcribed by neurons compared to 2 – 4% by RPE cells, which was the main *cfh* producer **(Fig 2F)**. The *cfi* transcript levels doubled in neurons and RPE cells of aging mice, while microglia from 24 week old mice produced less than 50% of the amount of those from 8 week old mice. Remarkably, *cfi* together with its functional counterpart *cfp* were the only complement transcripts that we tested and which were upregulated in neurons of 24 week old mice **(Fig 2F)**.

In summary, our data revealed an age-related change in cell type-specific complement expression towards a pro-inflammatory milieu, as complement activating molecules are upregulated and the *cfh* transcript amount was reduced across the entire retina.

### A characteristic proportion of activating and inhibiting complement transcripts in distinct retinal cell types

In our proof-of-concept experiments, we showed that five different retinal cell populations exhibited a cell type-specific expression of *c1s, c3, cfb, cfp, cfh* and *cfi* in albino BALB/c mice **(Fig 2F)** and thus may modulate cell type-specific retinal complement activity. Next, we investigated the balance of complement activator and inhibitor expression in the different cell types in more detail, this time normalizing expression levels to our novel housekeeping genes to allow comparability between cell populations independent of their cell counts in the retina. Strikingly, by this approach we distinguished cell types expressing mainly inhibiting complement components (*cfh, cfi*), like RPE and microglia, from cells, which synthetized mostly complement activating (*c1s, c3, cfb, cfp*) mRNA (neurons and Müller cells) in albino mice **(Fig 3A)**. We found that neurons and Müller cells mainly expressed *cfp*, a positive regulator of the complement system, while the remaining populations primarily synthetized transcripts of *cfh*, known to reduce complement activity **(Fig 3A)**. Interestingly, the most frequent complement negative regulator produced by all retinal cell types was *cfh* except for neurons which expressed more *cfi* than *cfh* mRNA **(Fig 3A).**

Taken together, our results demonstrate that the complement expression pattern of Müller cells and neurons of albino mice is clearly distinguishable from that of RPE, vascular and microglia cells.

### Early cell type-specific changes in the retinal complement expression contribute to hereditary retinal degeneration

We showed that the cellular complement expression profile **(Fig 3A)** and the retinal complement composition is altered during aging in albino mice **(Fig 2F)**. From this, we suggest that an early retinal cell type-specific shift in complement expression may contribute to retinal degeneration. To follow up this hypothesis, we used albino mice lacking the functional ABCA4 transporter (ABCA4^-/-^ mice) as a model for slow hereditary retinal degeneration ^45^. In these mice, the retinae showed a reduced number of cell nuclei in the ganglion cell layer and outer nuclear layer, accompanied by an enhanced autofluorescence in the RPE of animals older than 32 weeks as compared to wild type controls **(Fig EV3A – D)**. Various studies demonstrated an enhanced complement activity in RPE cells and whole eye extracts in these mice and it has been speculated about an involvement of the complement system in this degenerative process ^29–31,46,47^. Here, we analysed the contribution of the five major retinal cell types to modulate the local complement homeostasis in the eye of aging ABCA4^-/-^ mice. We found that in 24 week-old ABCA4^-/-^ mice complement-activating *c3* transcripts were stronger upregulated in RPE cells but reduced in neurons as compared to wild type littermates **(Fig 3B)**. Additionally, we detected more mRNA of the complement amplifier *cfp* in microglia cells of the ABCA4^-/-^ retina than in age-matched 24 week old wild type controls **(Fig 3B)**. Transcripts of the opposing complement-inhibiting *cfi* were downregulated in aged ABCA4^-/-^ microglia as compared to microglia from wild type albino mice (**Fig 3B**). These changes in the local complement expression pattern towards pro-inflammation in ABCA4^-/-^ mice at 24 weeks of age compared to wild type mice were present at a time point where retinal cell loss was first detected in the ganglion cell layer and, thus, occurred concomitantly (**Fig 3B, EV3A-C**). The enhanced complement activity was associated with a modified immune homeostasis in ABCA4^-/-^ retinae indicated by increased numbers of microglia that display signs of mild activation such as shorter microglial processes (**Fig EV3E**). These findings are well in line with the enhanced expression of activators (*c3, cfb, cfp*) and a downregulation of inhibitors (e.g. *cfi*) of the complement system in the retinal microglia (**Fig 3B**).

In conclusion, our data imply a close interplay of changes in the cell type-specific retinal complement component expression during hereditary retinal degeneration in ABCA4^-/-^ mice and activation of microglia cells that possibly contribute to slow neurodegeneration.

### Differences in complement expression in human, albino and pigmented murine retinae

It should be emphasized that the findings in albino mice did not generalize to murine or even human retinae with normal RPE pigmentation **(Fig 4A, Fig EV4)**. Similar to what we found in albino mice, Müller cells and neurons of pigmented C57BL/6 mice primarily expressed transcripts of complement activators, but microglia and vascular cells showed a different complement expression profile than in age-matched albino mice **(Fig 4A)**. We detected more complement activating components, especially *cfb*, in microglia and vascular cells of C57BL/6 mice than in the BALB/c mice **(Fig 4A)**. In RPE cells the ratio of complement activating and inhibiting transcripts was comparable between albino and pigmented mice.

Human retinal cells expressed a different ratio of complement inhibitors and activators compared to mouse retinal cells **(Fig 3A, 4A, EV4)**. In contrast to murine retinal cells, we observed a predominant transcript expression of *c1s* in RPE and neurons from the human retina and rather low transcript levels of *cfp* as determined by RNA sequencing **(Fig EV4)**. These interspecies differences in the expression of complement components may result in a more active complement system in human RPE and retinal neurons compared to their murine counterparts. In human Müller cells, local complement inhibition appeared to dominate since more than 75% of the tested transcripts were those for the complement inhibitors *cfi* and *cfh* **(Fig EV4)**. On the basis of our transcriptome data, we conclude that *cfi* could be the main complement negative regulator in the human retina but not *cfh* as shown before for murine retina **(Fig EV4)**.

In summary, our data imply a characteristic balance of activating and inhibiting complement transcripts in the different retinal cell types. Importantly, this cell type-specific expression of complement transcripts varied between pigmented and non-pigmented mouse strains and considerably between murine and human retinal cells.

### Acute ischemic retinal damage elicits a robust intraretinal, cell type-specific complement activation

We described an altered cell type-specific complement expression in the aging and degenerating retina of albino mice **(Fig 2, 3)**. In contrast, pigmented mice differed in their complement expression profile compared to albino mice **(Fig 4A)**. The majority of the analysed transcripts in the isolated cell types of pigmented retinae were that of complement activators (64-88%), whereas untreated RPE cells showed a balanced expression between analysed complement activators and inhibitors **(Fig 4A).** To evaluate changes in the cell type-specific complement expression in pigmented murine retinae upon tissue damage, we used a retinal ischemia/reperfusion (I/R) injury model to induce acute retinal degeneration ^48^ **(Fig 4)**. We found a massive rise in activating complement component expression 24 hours post ischemia in the different isolated cell populations that was more pronounced than described in aging or ABCA4^-/-^ mice **(Fig 2F, 3B, 4)**. The expression rates of complement activating transcripts *c1s, c3* and *cfb* summed up over all retinal cell populations were highly upregulated by 29%, 80% and 67% in post ischemic retinae **(Fig 4C)**. Interestingly, this was mainly mediated by expression changes in the RPE **(Fig 4C)**. The overall transcript amount of the only known positive regulator of the complement system, *cfp*, more than doubled in RPE and rose about 25% when considering the whole retinal transcript level **(Fig 4C)**. The analysed inhibitory complement regulators *cfh* and *cfi* showed a differential pattern of expression regulation **(Fig 4B)**. While *cfh* mRNA amounts were reduced, those of *cfi* were upregulated in cells from I/R retinae **(Fig 4C)**. Interestingly, *cfh* transcripts were provided mainly by the Müller cell and RPE populations whereas the neuronal population rather exclusively expressed *cfi* **(Fig 4C)**. This was consistent with the normalized expression ratios of complement components in the different cell populations for which we identified *cfi* as the main complement inhibitor in neurons but *cfh* as dominant complement inhibitor in the remaining cell populations **(Fig 4A)**.

Our results imply a key role of retinal cell types putatively shaping the initial immune response leading to retinal degeneration by expressing complement components and thereby priming complement activation 24 hours following retinal I/R injury.

## Discussion

### Major contributions of Müller cells and microglia to the retinal transcriptome

The retina consists of more than 40 distinct cell populations ^1,49^. So far, almost all transcriptomic analyses used whole retinae, therefore yielding RNA profiles averaging expression profiles from a mixture of cells ^5^. We enriched the five major retinal cell populations that significantly contribute to normal retinal function: Müller cells, microglia, vascular cells, retinal neurons and RPE cells ^42^. Our results demonstrated that previous studies possibly underestimated the contribution of Müller cells and microglia to the retinal transcriptome. We show a 7 – 9-fold higher proportional contribution of Müller cells and 5 – 60-fold higher contribution of microglia to the total retinal transcriptome than compared to other studies where their contribution was simply estimated on the basis of cell numbers ^43^. Quantification of cell numbers does not allow to draw conclusions about the contribution of the respective cellular entity to the global expression profile of the retina. Obviously, the overall transcriptional activity and therefore the potential influence of the different cell populations on retinal physiology could be independent of their cell counts. Transcription is highly regulated and it is known that cells with larger cell bodies such as Müller cells can provide more mRNA compared to cells with a smaller cell volume but with a higher cell number (e.g. neurons, RPE) ^50,51^.

### Defined cell types locally express and regulate complement transcripts within the retina

The complement system is involved in the maintenance of ocular functions ^52–54^ and in retinal pathologies ^11,12,29,39,46,55,56^. Compelling evidence suggests that complement components are locally produced in the retina ^15–17^. Mainly microglia ^23,24^ and RPE/choroid ^15,16,25^ had been described as a source of local complement production. Using cell type-specific complement expression analysis, we now show for the first time that at least five different retinal cell populations are capable to modulate retinal complement homeostasis. These cell populations expressed *c1s, c3, cfb, cfp, cfh* and *cfi* at varying levels depending on age, species, pigmentation and retinal degeneration. This corresponds to previous work describing complement transcripts in human ^2,17,18^, mouse ^15,16,19–22^ and rat retinae ^23^ without specifying the producing cell types within the tissue. Quantitative measurements of mRNA transcripts do not necessarily reflect rates of translation, and protein turnover from the different cellular identities. Nevertheless, we found complement proteins attached to isolated and properly washed cell populations further supporting the concept of a local production of complement proteins in the eye. Importantly, quantitative mRNA analysis can provide a molecular snapshot of the cellular response to changes in the retinal microenvironment, e.g. caused by accumulation of oxidative stress epitopes or deposition of cellular debris, proteins and lipids as seen in retinal drusen, especially during retinal degeneration like in AMD.

The regulation of complement expression in whole cell populations from the aging albino retina largely matched the changes we calculated for the normalised cellular expression rates in the distinct cell types. This implies that expression changes in most cases did not account for major changes in cell numbers, but rather appeared to be due to changes at transcriptional level. However, we cannot exclude the possibility of interfering effects. As an example, we found that in vascular cells the transcript amount of *c3, cfb* and *c1s* increased during aging when normalized to the housekeeping gene. This effect was diminished when we evaluated the overall expression of these complement components in the whole vascular cell population, which could possibly be explained by loss of vascular cells with the progress of aging. Additionally, the relative expression of all tested complement components decreased in RPE with increasing age except for *cfh*, which was expressed more strongly. In contrast, the absolute *cfh* transcript level detected in the whole RPE population of 24 week old mice was reduced. We speculate that the *cfh* upregulation in the surviving RPE of the aging eye could not counterbalance the overall age-associate RPE cell loss or dysfunction (as also indicated by a reduced *rpe65* expression). Accordingly, the RPE-dependent *cfh* transcript amount decreased and, thus, its putative contribution to balance the complement activity in the retinal microenvironment was diminished.

We found that the majority of the murine and human *cfh* transcript is provided by the RPE, which is in accordance with the work of Li et al. ^18^ reporting higher levels of *cfh* in the human RPE than in den retina. We identified neurons in the murine retina as the major producers of additional complement regulators such as *cfp* and *cfi,* which can either counteract or facilitate *cfh* function, respectively. The putative relevance of these complement components for retinal pathomechanisms was indicated by polymorphisms in the *cfi* and *cfh* genes being associated with an altered risk to develop AMD ^11,57^. Moreover, accumulation of CFP in deposits of AMD patients has been documented ^58^. In murine models of retinal degeneration, *cfi* expression was increased after polyethylene glycol induced insult and *cfp* expression was decreased in the light-damaged retina ^15,59^. We provide first evidence that *cfi* and *cfp* are primarily expressed by neurons that to date were not considered to contribute to the modulation of the retinal immune response. Accordingly, by expressing these complement components, they can potentially affect disease progression in damaged murine retina ^15,59^ or in AMD patients ^57,58^.

In our study, human retinal cells display a different cell type-specific complement expression profile than those of mice. We identified *cfi* as the putative main complement inhibitor in the human retina in contrast to *cfh* dominating in the murine retina which correlates with previous data from whole retinal transcript profiles ^60^. Additionally, the relative *c1s* expression was higher than that of most other complement activating components in human than in murine retinal tissue. Differences between the human and murine complement system have already been described by others ^61^: some human complement components are missing (e.g. factor H-related proteins), while others are duplicated (DAF, CD59) in the mouse. Some complement components can only be found in the murine complement system (CRRY). Given the differences in the complement expression profile of murine and human retinal cell types characterized in the present study and other variations in the murine and human retina (e.g. different cellular composition especially regarding photoreceptors, absence of a macula in the rodent retina), our findings emphasize that generalizing data regarding the contribution of the complement system to retinal pathology is difficult. Consequently, this also implies that translation of novel therapeutic strategies targeting the retinal complement system from mouse to human has to be done with caution.

### Age-dependent changes in cell type-specific complement expression

Age-related anatomical alterations in the retina are known from histological analyses ^62^. In albino retinae, we measured increased microglial and decreased RPE marker gene expression in the course of aging. We confirmed previous results showing an increased number of microglia and a putative RPE loss in immunostainings during aging ^63–65^. Local complement activity is important for healthy aging of the retina as complement deficiency in mice results in decreased amplitudes of electroretinograms and a thinner inner nuclear layer in 6 – 12 month old mice ^66,67^. In our study, the expression of the complement transcripts *c1s, cfb, cfp* and *cfi* increased while that of *cfh* mRNA decreased in retinal cells of albino mice from 8 to 24 weeks of age. This may indicate a retinal adaptation to age-related accumulation of factors leading to cellular stress and ultimately to degenerative processes. Age-dependent upregulation of complement transcripts, including *c1q, c3, c4* and *cfb* in the retina was described by Chen et al., but the contribution of distinct cell populations remained unsolved ^26^. Our findings indicate a role of microglia to an enhanced complement activity as they increased the expression of *c3* and *cfb* during aging in albino mice. This is in line with findings previously reported for microglia purified from retinae of pigmented mice ^27^. Upregulation of complement components was previously also associated with senescent RPE cells and a concomitantly altered immune regulation has been described ^28^. However, the contribution of Müller cells, retinal neurons and vascular cells in age-dependent complement expression so far was unknown. Here we present an original data set demonstrating that Müller cells and neurons provide a major percentage of retinal complement component transcripts and, thus, very likely have a major and to date largely underestimated impact on retinal complement homeostasis. While *c1s, cfp* and *cfi* expression significantly increased in Müller cells of older mice, neurons increased their *cfi* expression, but decreased that of *c1s* in 24-week-old albino mice. It is known from cell culture studies that Müller glia can produce C1q ^68^ and that complement activation products can regulate Müller cells via C5a-receptor contributing to retinal diseases ^69^. Our study suggests a direct involvement of Müller cells in the regulation of the retinal complement pathway even when the blood-retinal barrier is intact. It is well known from other parts of the central nervous system (CNS) that neurons and glia cells can provide complement components, like C1q, to orchestrate mechanisms during brain development (e.g. synaptic pruning, progenitor proliferation, neuronal migration) ^70,71^ and in the course of disease progression ^72,73^. Aging and Alzheimer diseased brains increase expression of *c1q, c3* and *c4* ^74,75^, which suggests a general mechanism of local complement function in the aging CNS and probably also in the retina.

### Modified complement expression in microglia and RPE of ABCA4-/-mice occurred before photoreceptor loss

Changes in retinal complement expression have been associated with a variety of retinal degenerative processes ^11,12,29,39,46,55,56^. Previous studies have discovered an altered complement homeostasis in ABCA4^-/-^ mice ^29–31,46,47^, which are used as a model for the inherited retinal dystrophy Stargardt disease type 1. Here, we determined the contribution of different retinal cell types to the local complement homeostasis in the eye of ABCA4^-/-^ mice and discovered a locally increased expression of *c3* in the RPE/choroid and decreased mRNA amounts of the complement inhibitor *cfi* in microglia. These changes in transcription were detected in the ABCA4^-/-^ retinae before significant photoreceptor loss was observed. Whole eye cup transcription analysis of 9-month old ABCA4^-/-^ mice showed significantly increased *c3* expression compared to wild type mice ^30^. Our study revealed the RPE as cell population increasing *c3* expression in ABCA^-/-^ compared to wild type mice. In line with our findings, others found an enhanced deposition of C3 cleavage products at the RPE ^29,31^. For the complement inhibitor *cfh* a 3-fold decreased expression in RPE of 4 week old ABCA4^-/-^ mice has been shown recently ^29^. Having investigated ABCA4^-/-^ mice with an age ranging from 8 to 24 weeks, we could not confirm these results, but rather found an up-regulation of *cfh* as an age-dependent adjustment of *cfh* expression in the RPE of albino ABCA^-/-^ mice. Concomitantly, a tendency towards an enhanced complement activity was detected in albino ABCA^-/-^ mice with increasing age, as the complement inhibitors *cfi* and *cfh* were downregulated in microglia and neurons, respectively. These findings correlate with that of others describing an increased detection of inflammation markers, e.g. C-reactive protein in the retina of ABCA4^-/-^ mice ^29^. Moreover, our results may imply that rebalancing the cell type-specific complement activity in inherited retinal diseases may help to prolong neuron survival.

### Complement involvement and post-ischemic retinal degeneration

Retinal I/R injury is a pathological hallmark of diabetic retinopathy and glaucoma. Gene profiling studies of whole mouse retinae implicate an important role of the complement pathway in I/R associated damages ^32^. Retinal *c1q, c1s, c1r, c2, c3 c4a* and *cfh* expression as well as protein deposition was time-dependently increased after transient ischemia in mice ^32–34,76^. Mostly, late post-ischemic time points (3 – 28 d) were chosen for analysis ^32,33,76^. Data by Kim et. al. suggest an early involvement of the complement system in retinal degeneration following I/R reporting an increase of retinal *c3* mRNA 24 hours after retinal detachment ^34^. However, it remains incompletely understood whether and how individual retinal cell types modulate the complement activity after retinal I/R injury. Here, we show that mainly Müller cells, microglia and the RPE respond with an altered complement expression upon ischemic tissue damage. *C3* expression was significantly enhanced in each of those cell types. Of note, we observed a concurrent post-ischemic upregulation of *cfb* expression in the same cellular populations. The positive complement regulator *cfp* was upregulated exclusively in micoglia. However, complement inhibitor *cfi*, showed cell type-specific enhanced mRNA levels in microglia, RPE and retinal neurons, whereas expression of the complement inhibitor *cfh* was significantly reduced in Müller cells. Consequently, these expression changes should lead to a boost of the complement pathway 24 hours after transient ischemia, indicating a potential early contribution to neurodegeneration.

As a consequence of energy deficiency, calcium imbalance and oxidative stress, retinal glia cells become activated after I/R injury ^48,77,78^. Importantly, microglia and Müller cells express a variety of complement receptors and a regulation of their physiological function by complement activation products has been demonstrated ^69,79,80^. Accordingly, changes of local complement expression, as observed in our study, holds the potential to influence the glial response to tissue damage.

Apart from its effects on glial functions, the complement system may also directly act on neurons. Retinal ganglion cells express the C5a receptor ^81^, indicating a susceptibility to C5activation products. Furthermore, complement-mediated retinal synapse elimination, caused by complement dependent synapse tagging, was implicated in retinal ganglion cell loss in glaucoma mouse models ^82^ and was proposed to be a general mechanism of complement activity in neurodegenerative diseases ^73^.

Even though not having investigated C5 expression, we demonstrated a strong activation of the complement system in general, with glia and RPE cells (and not neurons) as a major source of complement activators that ultimately should also induce enhanced formation of C5- activation products in the post-ischemic retina. The latter will likely promote retinal ganglion cell death. Findings from recent studies suggest that a reduced glial activation improves neuronal survival ^77,83^ and that a deficiency in C3 or C1q alleviated cell death after I/R ^33,76,84^. Together with our data this implies a key role of local modulations of the retinal complement homeostasis on the glial tissue response and, consequently, indirectly on neuronal survival. This intriguing hypothesis however needs further testing, ideally including expression analysis of additional relevant complement components like C5 or C9.

### Different cell types are involved in an altered complement expression in inherited and acute retinal degeneration

Our data revealed distinct complement expression profiles of the respective cell populations pointing at their different roles in determining the complement activity of the retinal microenvironment, depending on the underlying pathology. In the ABCA4^-/-^ albino mouse model of slow retinal degeneration, mainly cell types producing complement inhibitors, like microglia and RPE cells, adapted to the stress situation, while in the acute model of transient ischemia also Müller cells were involved in the modulation of complement homeostasis. In total, these three cell populations produced up to 95% complement activating transcripts determining the complement activity in the post-ischemic retina. The overall changes in complement expression were comparable to findings from previous microarray studies in whole post-ischemic retinae ^32^.

Genetic variation in several complement genes and specifically those of the regulators CFH and CFI are associated with AMD ^11,57,85^. CFI regulates the complement activity by degrading complement components C3b and C4b in the presence of specific cofactors, with CFH being its main cofactor ^86,87^, thereby facilitating the cleavage of C3b into inactive fragments ^88^. Surprisingly, we report a spatially distinct transcription pattern of *cfi* and its cofactor *cfh* in the murine and human eye. *Cfi* mRNA was mainly detected in retinal neurons of murine retinae, whereas in the human retina most *cfi* transcripts were present in Müller cells, too. However, *cfh* transcripts were primarily present in RPE cells both in mice and human retinae. Further, we found a counteractive transcriptional regulation of *cfi* and *cfh* during aging and in the context of retinal degeneration. *Cfi* transcripts were increased in the aging and post-ischemic murine retina, while *cfh* transcripts were decreased. These astounding findings suggest a CFH-independent function of CFI in the retina and could imply other relevant cofactors of CFI such as CR1 or CD46 ^88^. To date, there are no known AMD-associated polymorphisms in the *cd46* or *cr1* genes reported, but knockout of *cd46* induced a dry AMD-related phenotype in the mouse ^89^ and underpinned an important function of CD46 (possibly in conjunction with CFI) in retinal physiology.

Diabetic retinopathy is associated with polymorphisms in the *cfb* and *cfh* gene ^13^. We identified Müller cells as major players in the post-ischemic mouse retina which shows hallmarks of diabetic retinopathy. Indeed, these cells downregulated *cfh* transcripts and upregulated *cfb*.

## Conclusion

Our in-depth cell type-specific expression analyses yield valuable insights into aspects of retinal physiology and pathology. We identified strikingly different complement expression profiles in Müller cells, microglia, vascular cells, retinal neurons and RPE cells. These distinct profiles potentially determine the capability of these cell populations to modulate complement activity in the retinal microenvironment, independent of the systemic levels of complement expression. Given the considerable number of expressed complement genes in the juvenile retina, we hypothesize that local complement expression is key to maintain retinal integrity under normal conditions. Moreover, our data imply that modulations in local complement expression, primarily in glia and RPE cells during aging, chronic and acute stress can facilitate retinal cell death and retinal degeneration independent of the systemic complement system. A reaction common to I/R damage, aging and hereditary retinal degeneration (ABCA4^-/-^ mice) appears to be the disinhibition of complement activity by intraretinal downregulation of the complement inhibitor CFH. Furthermore, our cell type-specific transcription analysis provides a novel perspective on how expression of genes identified by gene association studies for AMD and diabetic retinopathy in various retinal cell types might explain the pathomechanisms in question. Finally, the seemingly fine-tuned reaction of all retinal cell types to distinct conditions of tissue stress suggests that this needs to be considered for successful development of novel therapeutic strategies targeting retinal complement activity.

## Materials and Methods

### Animals

All experiments were done in accordance with the European Community Council Directive 2010/63/EU and the ARVO Statement for the Use of Animals in Ophthalmic and Vision Research and were approved by the local authorities (55.2 DMS-2532-2-182). All mice were housed in a 12-hours light/ dark cycle with approx. 400 lux. Experiments were conducted with 8, 16 and 24-week-old male and female B albino *abca4*^-/-^ and *abca4*^+/+^ mice on BALB/c background ^90^, which were obtained by breeding of kindly provided breeding pairs by T. Krohne (University of Bonn, Germany). Genotyping of the mice was performed with the KAPA Mouse Genotyping Hot Start Kit (Merck, Darmstadt, Germany) using *abca4_wt_f* GCCCGCACTTGTGTATTTAG, *abca4_wt_r* GCCTTTTCCTCAGAGTCCAG, *abca4_ko_f* TGAATGAACTGCAGGACGAG and *abca4_ko_r* AATATCACGGGTAGCCAACG primer. Retinal ischemia was induced in 8 weeks old C57BL/6J mice as previously described ^77,78^. Briefly, transient retinal ischemia was induced in one eye by the high intraocular pressure (HIOP) method. The other eye remained untreated as internal control. Anesthesia was induced with ketamine (100 mg/kg body weight, intraperitoneal (ip); Ratiopharm, Ulm, Germany), xylazine (5 mg/kg, ip; Bayer Vital, Leverkusen, Germany), and atropine sulfate (100 mg/kg, ip; Braun, Melsungen, Germany). The anterior chamber of the test eye was cannulated from the pars plana with a 30-gauge infusion needle, connected to a saline bottle. The intraocular pressure was increased to 160 mmHg for 90 minutes by elevating the bottle. After removing the needle, the animals survived for 24 hours and, subsequently, were sacrificed with carbon dioxide for tissue analysis.

### Human donor eyes

Eye samples of donors were provided by Prof. Dr. Margaret DeAngelis (Department of Ophthalmology and Visual Sciences, John Moran Eye Center, University of Utah School of Medicine, Salt Lake City, Utah, USA) who received the eyes from the Utah Lions Eye Bank (Salt Lake City, Utah, USA). Post-mortem time ranged between 4 and 6 hours. The study was performed in accordance with the tenets of the Declaration of Helsinki and the Medical Research Involving Human Subjects Act (WMO) and was approved by the local ethics committee of the University of Utah, School of Medicine (Salt Lake City, Utah, USA). Informed consent from all deceased individuals’ family donors for tissue donation was obtained.

### Isolation of retinal cell populations by immunomagnetic enrichment

Retinal cell types were enriched as described previously ^42^. Briefly, retinae were treated with papain (0.2 mg/ml; Roche Molecular Biochemicals) for 30 minutes at 37 °C in the dark in Ca^2+^-and Mg^2+^-free extracellular solution (140 mM NaCl, 3 mM KCl, 10 mM HEPES, 11 mM glucose, pH 7.4). After several washes and 4 minutes incubation with DNase I (200 U/ml), retinae were triturated in extracellular solution (now with 1 mM MgCl_2_ and 2 mM CaCl_2_). To purify microglial and vascular cells, the retinal cell suspension was subsequently incubated with CD11b-and CD31 microbeads according to the manufactures protocol (Miltenyi Biotec, Bergisch Gladbach, Germany) and the respective binding cells were depleted from the retinal suspension using LS-columns prior to Müller cell enrichment. To purify Müller glia, the cell suspension was incubated in extracellular solution containing biotinylated hamster anti-CD29 (clone Ha2/5, 0.1 mg/ml, BD Biosciences, Heidelberg, Germany) for 15 minutes at 4°C. Cells were washed in extracellular solution, spun down, resuspended in the presence of anti-biotin MicroBeads (1:5; Miltenyi Biotec,) and incubated for 10 minutes at 4°C. After washing, CD29+ Müller cells were separated using large cell (LS) columns according to the manufacturer’s instructions (Miltenyi Biotec). Cells in the flow through of the last sorting step were considered as neuronal population as it was now depleted from microglia, vascular cells and Müller glia. Retinal pigment epithelium was collected by scratching it out of the eye cup after the retina had carefully been removed and, thus, scratch samples also contained cells from the underlying choroid. In some experiments, RPE scratch samples were depleted from choroidal macrophages and vascular cells. Samples were digested and in subsequent steps macrophages were depleted using anti-CD11b-microbeads and vascular cells using CD31-microbeads (Miltenyi Biotec) like described above for retinal samples.

### RNA sequencing

Total RNA was isolated from cell pellets after immunoseparation using the PureLink^®^ RNA Micro Scale Kit (Thermo Fisher Scientific, Schwerte, Germany). RNA integrity validation and quantification was performed using the Agilent RNA 6000 Pico chip analysis according to the manufactures instructions (Agilent Technologies, Waldbronn, Germany). Enrichment of mRNA and library preparation (Nextera XT, Clontech), library quantification (KAPA Library Quantification Kit Illumina, Kapa Biosystems, Inc., Woburn, MA, USA) as well as sequencing on an Illumina platform (NextSeq 500 High Output Kit v2; 150 cycles) were performed at the service facility of the KFB Center of Excellence for Fluorescent Bioanalytics (Regensburg, Germany; www.kfb-regensburg.de). After de-multiplexing, at least 20 million reads per sample was reached. Quality control of the reads and quantification of transcript abundance was performed with the Tuxedo suit, as described elsewhere ^91^. To this end, *cutadapt* was used to remove adapter sequences ^92^ and several quality control measures were queried with *fastqc*. No major problems with the sequencing data was detected. Next, the trimmed reads were aligned to the reference genome/transcriptome (*mm10*) with *HISAT2* ^93^ and transcript abundance was estimated with *stringtie,* expressed as fragments per 1,000 base pairs of transcript per million reads (FPKM).

### qRT-PCR

Like for RNA seq total RNA was isolated from enriched cell populations using the PureLink^®^ RNA Micro Scale Kit (Thermo Fisher Scientific, Schwerte, Germany). A DNase digestion step was included to remove genomic DNA (Roche). In addition, RNA integrity validation and, importantly, quantification was performed using the Agilent RNA 6000 Pico chip analysis according to the manufactures instructions (Agilent Technologies, Waldbronn, Germany). First-strand cDNAs from total RNA purified from each cell population were synthesized using the RevertAid H Minus First-Strand cDNA Synthesis Kit (Fermentas by Thermo Fisher Scientific, Schwerte, Germany). Primers were desgined using the Universal ProbeLibrary Assay Design Center (Roche). Transcript levels of candidate genes were measured by qRT-PCR using cDNA with the TaqMan hPSC Scorecard(tm) Panel (384 well, ViiA7, Life Technologies, Darmstadt, Germany) according to the company’s guidelines.

### LC-MS/MS mass spectrometric analysis

LC-MS/MS analysis was performed as described previously ^94,95^ on an Q Exactive HF mass spectrometer (Thermo Fisher Scientific Inc., Waltham, MA, U.S.A.) coupled to a Ultimate 3000 RSLC nano-HPLC (Dionex, Sunnyvale, CA). Approximately 0.5 μg per sample were automatically loaded onto a nano trap column (300 μm inner diameter × 5 mm, packed with Acclaim PepMap100 C18. 5 μm, 100 Å; LC Packings, Sunnyvale, CA) before separation by reversed phase chromatography (HSS-T3 M-class column, 25 cm, Waters) in a 80 minutes non-linear gradient from 3 to 40% acetonitrile (ACN) in 0.1% formic acid (FA) at a flow rate of 250 nl/min. Eluted peptides were analysed by the Q-Exactive HF mass spectrometer equipped with a nano-flex ionization source. Full scan MS spectra (from m/z 300 to 1500) and MSMS fragment spectra were acquired in the Orbitrap with a resolution of 60,000 or 15000 respectively, with maximum injection times of 50 ms each. The up to ten most intense ions were selected for HCD fragmentation depending on signal intensity (TOP10 method). Target peptides already selected for MS/MS were dynamically excluded for 30 seconds. Spectra were analyzed using Progenesis QI software for proteomics (Version 3.0, Nonlinear Dynamics, Waters, Newcastle upon Tyne, U.K.) for label-free quantification as previously described^44^. All features were exported as Mascot generic file (mgf) and used for peptide identification with Mascot (version 2.4) in the UniProtKB/Swiss-Prot taxonomy mouse database (Release 2017.02, 16 871 sequences). Search parameters used were: 10 ppm peptide mass tolerance and 20mmu fragment mass tolerance, one missed cleavage allowed, carbamidomethylation was set as fixed modification, methionine oxidation and asparagine or glutamine deamidation were allowed as variable modifications. A Mascot-integrated decoy database search calculated an average false discovery of < 1%.

### Western Blot

Cell pellets of enriched cell populations from pooled two mouse eyes were dissolved in reducing Laemmli sample buffer, denatured and sonicated. Neuronal protein extraction reagent (Thermo Fisher Scientific, Braunschweig, Germany) was added to the neuron populations. Samples were separated on a 12% SDS-PAGE. The immunoblot was performed as previously described ^15^. Detection was performed with primary and respective secondary antibodies diluted in blocking solution **(Appendix Table S1)**. Blots were developed with WesternSure PREMIUM Chemiluminescent Substrate (LI-COR, Bad Homburg, Germany).

### Immunohistochemistry of retina and RPE/choroid flat mounts

To quantify cell nuclei in retinal layers sections of 4% paraformaldehyde (PFA)-fixated and paraffin-embedded murine eyes were deparaffinised and incubated with Hoechst33342 (1:1000; #H1399, Thermo Fisher Scientific, Braunschweig, Germany) as previously described ^15^. Images were acquired using confocal microscopy (VisiScope, Visitron Systems, Puchheim, Germany).

Retinal microglia and RPE autofluorescence quantification was performed in retinal or RPE/choroid flat mounts, respectively. Anterior segments of mouse eyes were removed. Retina and RPE/choroid were carefully separated. Flat mounts were fixated in 4% PFA (RPE/choroid 2 h on ice, retina 1 h room temperature), permeabilized (1% Triton X-100) and blocked (1% BSA, 5% goat serum, 0.1 M NaPO_4_, pH 7). RPE/choroid flat mounts were stained with anti-ZO-1 antibody (over night, 4° C) and secondary antibody including Hoechst33342 (1 h, room temperature) **(Appendix Table S1)**. Retinal flat mounts were stained with anti-Iba1 antibody (3% Triton X-100, 1 % DMSO, 5% normal goat serum, over night 4° C) and secondary antibody (1% BSA in PBS, over night 4° C). Retinal flat mounts were embedded with photoreceptors down and ganglion cell layer facing up. Images were taken with a confocal microscope (VisiScope, Visitron Systems).

## Acknowledgements

This project was supported by the Deutsche Forschungsgemeinschaft (DFG-GR 4403/5-1 to AG and DFG-PA 1844/3-1 to DP). We thank Gabriele Jäger, Dirkje Felder, Renate Foeckler, Andrea Danullis and Elfriede Eckert for excellent technical support of cell preparation, immunodetections and molecular biology. Furthermore we thank Prof. Tim Krohne (University Hospital of Bonn, Germany) providing the ABCA4^-/-^ mice and Margaret DeAngelis (Department of Ophthalmology and Visual Sciences, John Moran Eye Center, University of Utah School of Medicine, Salt Lake City, Utah, USA) for supply with human donor eyes.

## Author contributions

DP, NS, SMH and AG designed research; DP, NS, FG, AP, TS, SMH and AG performed research; DP, NS, SMH, TS, BHFW, SMH and AG analysed and interpreted the data; DP and AG wrote the manuscript. All authors provided input to edit the manuscript.

## Conflict of interest

All authors declare no conflict of interest.

## The paper explained

### Problem

Activation of the complement cascade, a pathway of the innate immune system, can be observed in severe vision loss, e.g. caused by age-related macular degeneration, diabetic retinopathy or inherited retinal degeneration. Despite its importance, the regulation of retinal complement activity is still far from understood. It has been hypothesized that retinal complement activation may be mediated either locally by microglia/macrophages and/or the retinal pigment epithelium (RPE) cells or systemically via the liver, regardless of the fact that an intact blood-retinal barrier prevents systemically circulating proteins from entering the immune privileged retinal tissue.

### Results

We describe the liver-independent retinal complement system in the mouse and human retina by cell type-specific complement analysis. We report that five different retinal cell populations express complement components. These results indicate a modulation of the retinal complement homeostasis, independent of the complement components present in the systemic blood circulation. Cell type-specific complement expression is different in human/mouse, pigmented/non-pigmented and healthy/damaged retinae. The ratio of complement inhibitor expression *cfi/cfh* is proportionally higher in human than murine retinae.

Increased expression of complement activators was a general observation in the damaged retina, with the complement inhibitor *cfh* being downregulated. In contrast, the other major complement inhibitor *cfi* was found upregulated in the damaged retinae.

### Impact

We have discovered that local, retinal expression of complement components during aging and retinal degeneration is cell type-specifically regulated. These findings may change the perspective on how the retinal complement system contributes to retinal degeneration. Consequently, this work may help to develop novel therapeutic strategies better suited to target the complement system in the context of age-related macular degeneration, diabetic retinopathy or inherited retinal degeneration. These strategies should aim at modulating cell type-specific complement expression and, thereby, reducing the detrimental effects of an overshooting innate immune response in the process of retinal degeneration.

---

**Tab 1.** Dissection of the molecular and cellular architecture of the murine retina – contribution of distinct retinal cell types

Tables and their legends
|  |  |  |  |  |
| --- | --- | --- | --- | --- |
|  | whole RNA | *idh3b* transcript | Drop-Seq. <sup>1</sup> | Microscopy<sup>43</sup> |
| Müller cells | 31% | 25% | 3.6% | 2% |
| Microglia | 12% | 5% | 0.2% | 1% |
| Vascular cells | 4% | 1% | 0.6% | 1% |
| Retinal neurons | 30% | 60% | 95.3% | 85% |
| RPE | 23% | 9% | n.d. | 13% |

Drop-Seq. droplet-based single cell RNA sequencing, n.d. not determined

**Expanded View Figure legends**

**Figure EV1.**
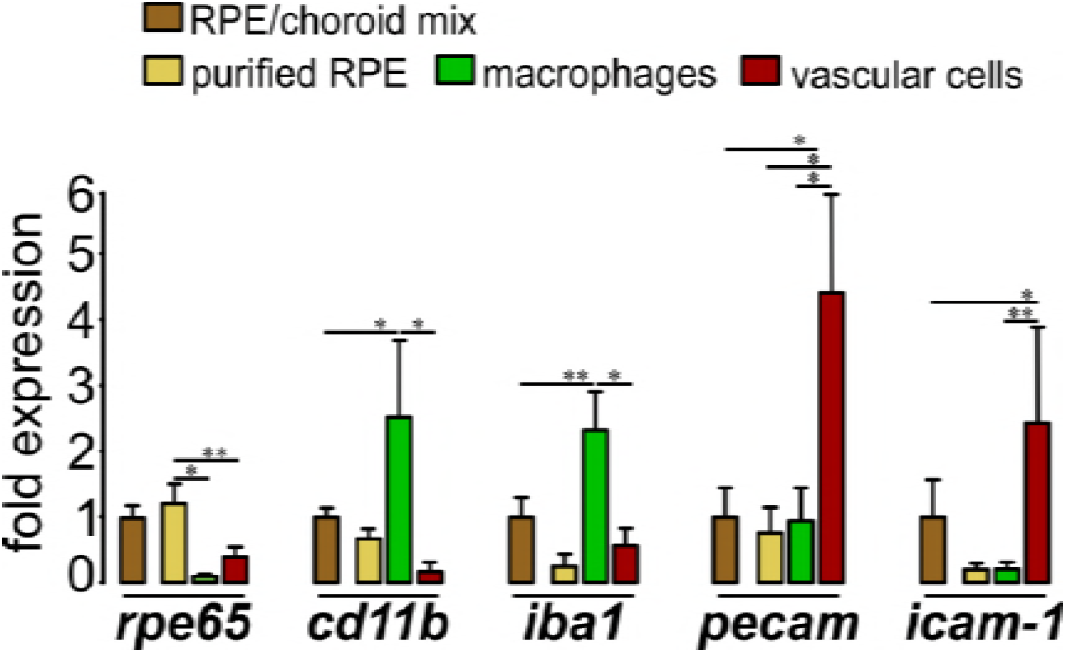
Characterization of different cell populations in the RPE/choroid scratch samples. Considerable expression of the macrophage marker genes *cd11b* and *iba1* as well as of *pecam* and *icam-1* as markers for vascular cells were detected in RPE/choroid scratch samples (RPE/choroid mix, brown). Implementing immunomagnetic depletion of CD11b^+^ macrophages (green) and pecam^+^ vascular cells (red) yielded purer RPE (yellow), indicated by reduced transcripts of *iba1* and *icam-1* in the purified RPE (yellow) compared to the RPE/choroid mixture (brown). Values show mean ± SEM (n=3 – 4, 8 weeks old C57BL/6; unpaired Student’s t-test: *P = 0.05, **P = 0.01)

**Figure EV2.**
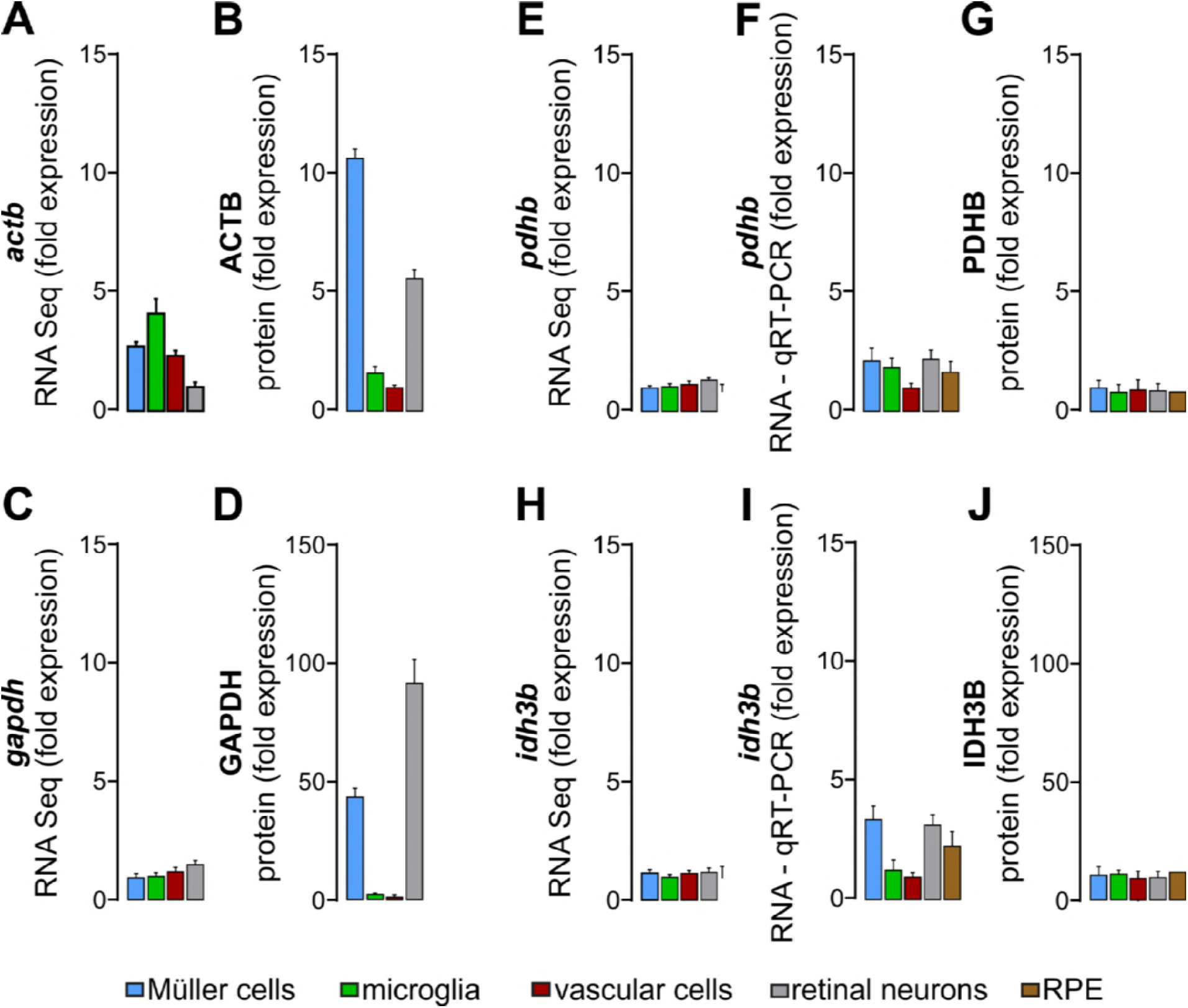
Putative housekeeping genes PDHB and IDH3b are expressed at comparable levels in all five investigated retinal cell populations. A, C, E, H RNA sequencing showed a similar expression levels of *gapdh, pdhb* and *idhb3* in Müller cells, microglia, vascular cells and retinal neurons, while *actb* expression showed considerable differences, e.g. between microglia and neurons (n=13, mean ± SEM) relative to total mRNA input amounts. B, D, G, J Quantitative mass spectrometric analysis revealed equal protein expression levels of PDHB and IDH3b relative to total protein input amounts in all investigated cell populations, while commonly used housekeeper like ACTB and GAPDH showed major differences regarding expression levels in the distinct cell populations (n=3 – 4, mean ± SEM). F, I Expression of housekeeping genes was comparable using qRT-PCR in all five cell populations for which input RNA had been exactly adjusted according to results from RNA quantification by RNA picochip 6000 analysis (n=3 – 4, mean ± SEM).

**Figure EV3.**
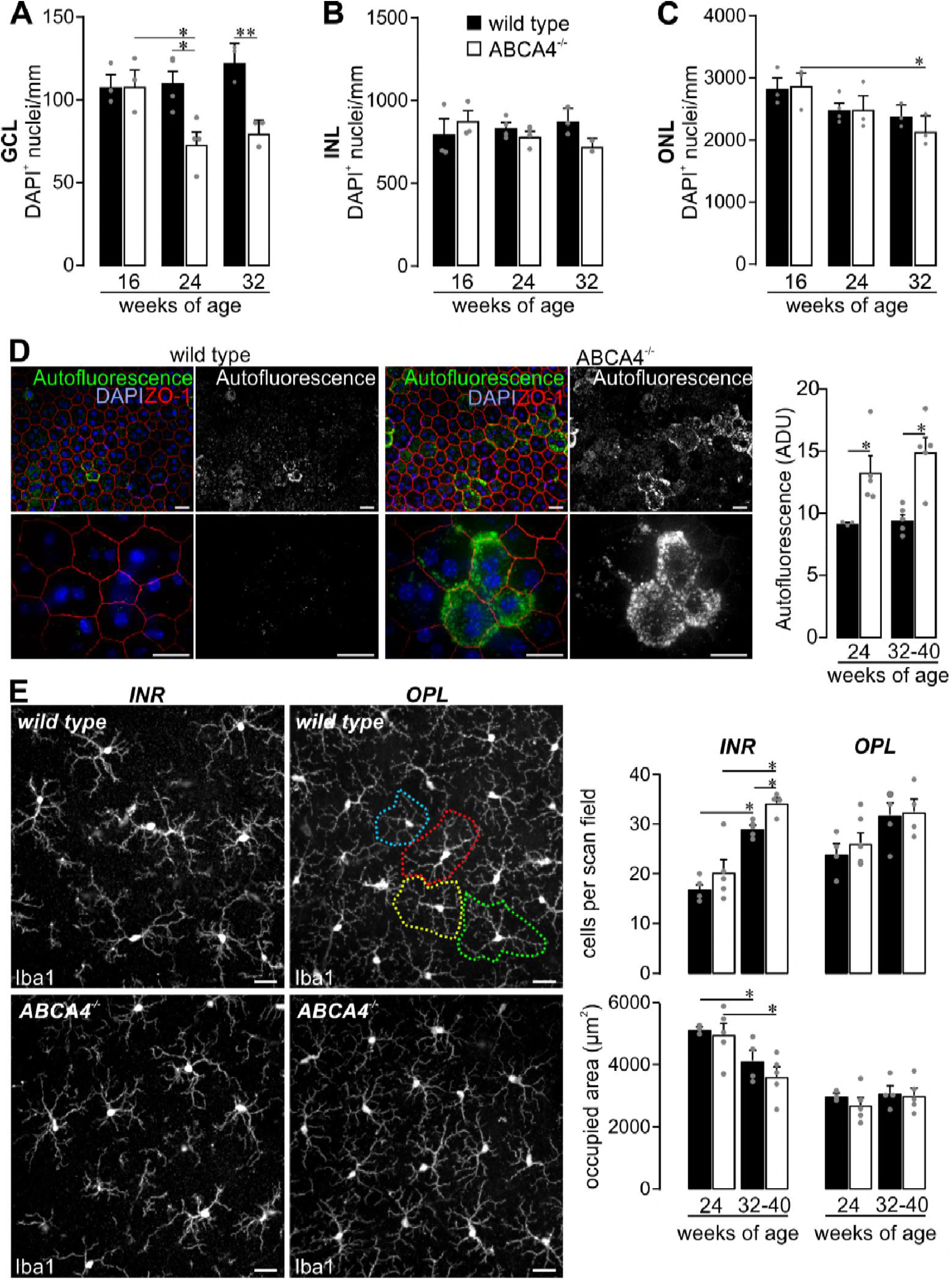
Retinal phenotype in the investigated albino ABCA4-/-strain. A The quantity of DAPI^+^ cell nuclei in the ganglion cell layer was decreased in 24 and 32 week old ABCA4^-/-^ mice but not in the wild type, albino retina. B Cell numbers in the retinal inner nuclear layer (INL) were stable in aging wild type and ABCA4^-/-^ mice. C Photoreceptor density was determined by quantification of DAPI^+^ cell nuclei in the outer nuclear layer (ONL). It significantly decreased in ABCA4^-/-^ mice at 32 weeks of age. D The integrity and autofluorescence of the RPE was investigated in fixed eye cup preparations after carefully removal of overlaying retinae. ZO-1 labelling (red) delineates tight junctions formed by intact RPE cells. Autofluorescence was detected upon excitation with a 488 nm laser and revealed enhanced signals from RPE of ABCA4^-/-^ mice as compared to RPE from wild type littermates. Mean grey values for autofluorescence were measured over the whole scan field. E *Right*, microglia were quantified in the inner retinal layers (INR) such as ganglion cell and inner plexiform layer and additionally in the outer plexiform layer (OPL) on basis of Iba1 labeling in mice of the indicated age. *Left*, the area occupied by a single microglia was measured as exemplarily depicted by the dashed circles of different colour for OPL microglia a retina from a wild type animal. A – E Bars represent mean values ± SEM from 2 – 4 animals. *P<0.05, **P<0.01, Mann-Whitney U-test. Scale bars, 20 µm. ADU, arbitrary digital units.

**Figure EV4.**
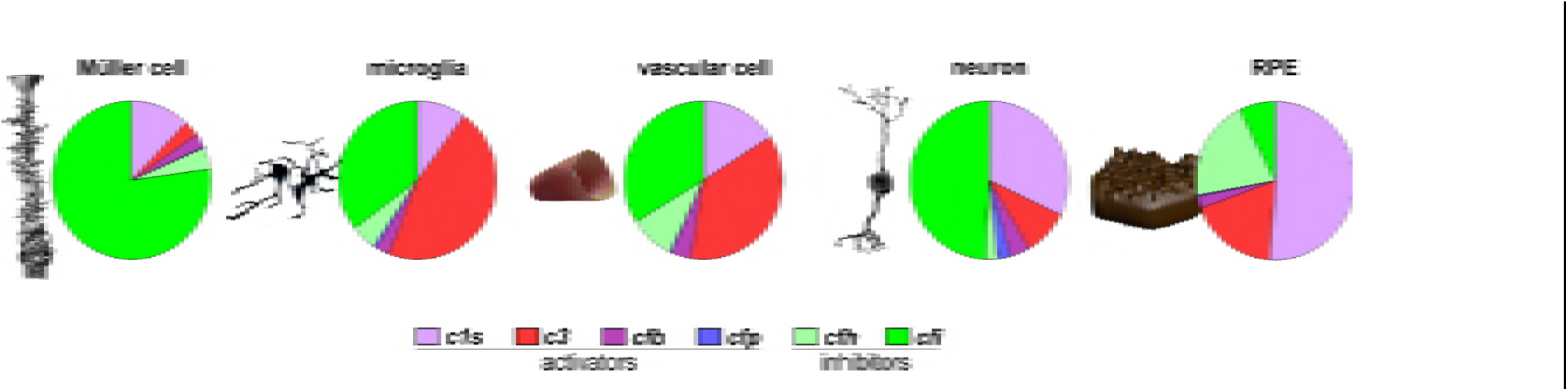
Complement expression profiles from cells of the human retina. The relative amounts of complement transcript expression per cell of the indicated complement components were compared in the five different human retinal cell populations using RNA sequencing analysis of four human retinae without obvious signs of retinal degeneration (numbers are given in **Table S4**).

## Funding

Deutsche Forschungsgemeinschaft DFG-GR 4403/5-1 to AG and DFG-PA 1844/3-1 to DP

**Appendix**

**Table S1.** Primary and secondary antibodies for Western Blot detection.

| Primary antibody | Species | Company | Catalogue number / Reference | Dilution / Concentration |
| --- | --- | --- | --- | --- |
| anti-PDHB | rabbit | Abcam (Cambridge, UK) | ab155996 | 1:1000 |
| anti-C1s | rabbit | Proteintech (Rosemont, IL, US A) | #14554-1-AP / <sup>15</sup> | 1:1000 |
| anti-C3-HRP | goat | MP Biomedicals (Santa Ana, C A, USA) | #55557 / <sup>15</sup> | 1:1000 |
| anti-CFB | goat | Merck (Darmstadt, Germany) | #341272 / <sup>15</sup> | 1:1000 |
| anti-CFP | rat | in-house | <sup>15</sup> | 1:250 |
| anti-CFH | goat | Merck (Darmstadt, Germany) | # 341276 | 1:500 |
| anti-CFI | goat | Quidel (San Diego, CA, USA) | A313 / <sup>96</sup> | 1:500 |
| anti-IBA1 | rabbit | Wako Chemicals (Neuss, Germany) | #019-19741 / <sup>97</sup> | 1:400 |
| anti-ZO-1 | rabbit | ThermoFisher (Braunschweig, Germany) | # 61-7300 / <sup>98</sup> | 1:300 |
| **Secondary antibody** |  |  |  |  |
| anti-rat Ig-HRP | goat | Dianova (Hamburg, Germany) | #112-035-003 | 1:5000 |
| anti-rabbit Ig-HRP | goat | Dianova (Hamburg, Germany) | #111-035-003 | 1:5000 |
| anti-goat Ig-HRP | rabbit | Dianova (Hamburg, Germany) | #305-035-003 | 1:5000 |
| Anti-rabbit-IG-Cy3 | goat | ThermoFisher (Braunschweig, Germany) | # A10520 | 1:200 |

**Table S2.** Percentage of murine, albino, aginig retinal cell type-specific complement expression profiles.

| Complement transcript | Müller cells |  |  | microglia |  |  | vascular cell |  |  | neuron |  |  | RPE |  |  |
| --- | --- | --- | --- | --- | --- | --- | --- | --- | --- | --- | --- | --- | --- | --- | --- |
|  | 8 | 16 | 24 | 8 | 16 | 24 | 8 | 16 | 24 | 8 | 16 | 24 | 8 | 16 | 24 |
| *c1s* | 9% | 5% | 11% | 2% | 3% | 2% | 4% | 14% | 11% | 8% | 7% | 2% | 9% | 4% | 4% |
| *c3* | 12% | 8% | 16% | 3% | 3% | 4% | 1% | 3% | 6% | 6% | 5% | 7% | 3% | 1% | 1% |
| *cfb* | 7% | 8% | 13% | 3% | 4% | 12% | 4% | 11% | 13% | 5% | 7% | 4% | 25% | 3% | 4% |
| *cfp* | 38% | 64% | 49% | 11% | 13% | 9% | 16% | 21% | 28% | 46% | 56% | 47% | 3% | 2% | 2% |
| *cfh* | 32% | 14% | 10% | 71% | 73% | 72% | 72% | 49% | 38% | 14% | 7% | 9% | 60% | 91% | 89% |
| *cfi* | 1% | 0.8% | 2% | 9% | 5% | 2% | 3% | 3% | 4% | 22% | 17% | 31% | 0.4% | 0.3% | 0.3% |

data for 8, 16 and 24 week old mice; graphically depicted in **Figure 3A**

**Table S3.**
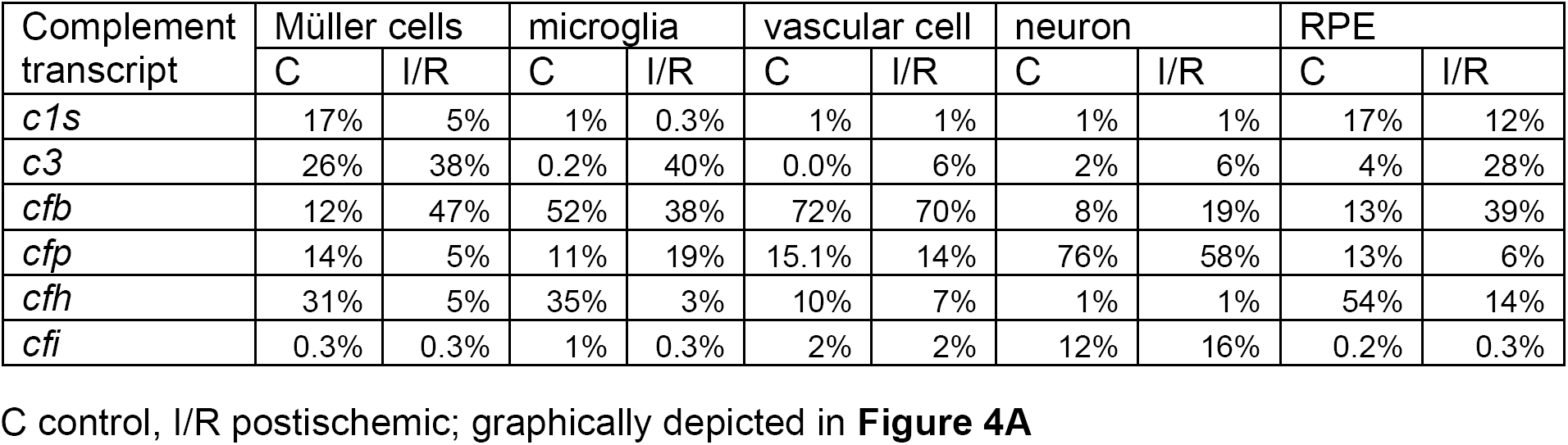
Percentage of murine, pigmented control and postischemic retinal cell type-specific complement expression profiles.

**Table S4.**
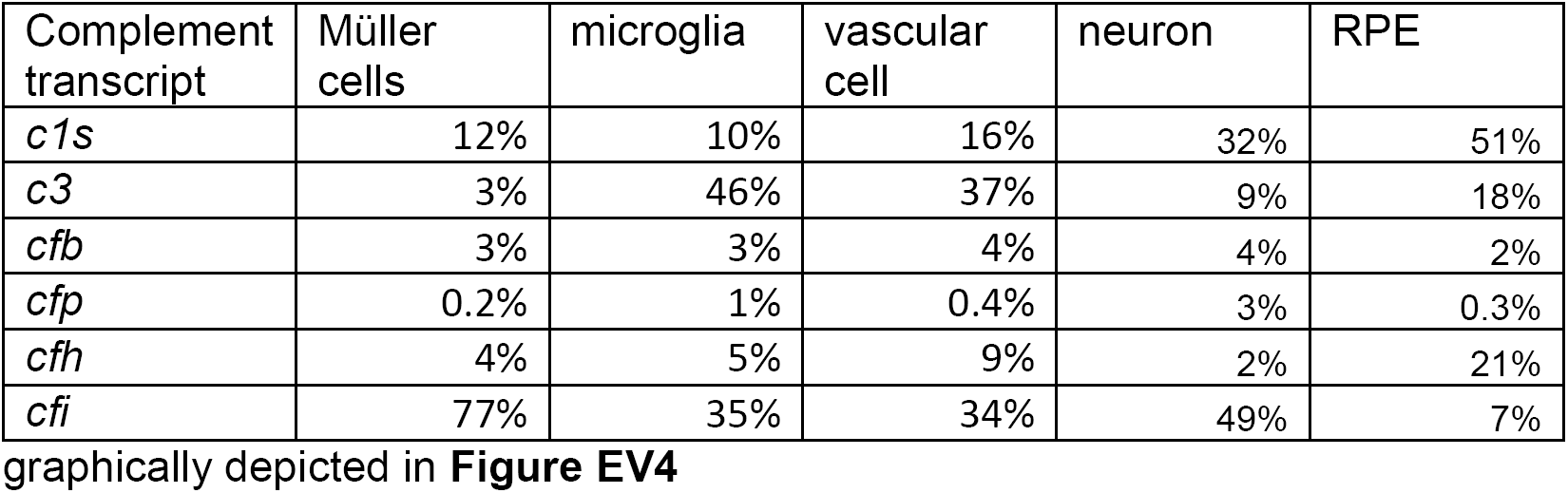
Percentage of human retinal cell type-specific complement expression profiles.

